# Purging of deleterious variants due to drift and founder effect in Italian populations with extended autozygosity

**DOI:** 10.1101/022947

**Authors:** Massimiliano Cocca, Marc Pybus, Pier Francesco Palamara, Erik Garrison, Michela Traglia, Cinzia F Sala, Sheila Uiivi, Yasin Memari, Anja Kolb-Kokocinski, Shane McCarthy, Richard Durbin, Paolo Gasparini, Daniela Toniolo, Nicole Soranzo, Vincenza Colonna

**Affiliations:** Department of Medical, Surgical and Health Sciences, University of Trieste, 34100 Trieste, Italy; Institut de Biologia Evolutiva (UPF-CSIC), Universitat Pompeu Fabra, Barcelona, Spain, 08003; Harvard T. H. Chan School of Public Health, 02115 Boston, MA.; Human Genetics, Wellcome Trust Sanger Institute, Genome Campus, Hinxton, CB10 1HH; Medical Genetics, Institute for Maternal and Child Health IRCCS “Burlo Garofolo”, 34100 Trieste, Italy; Division of Genetics and Cell Biology, San Raffaele Scientific Institute, Milano,Italy; Department of Haematology, University of Cambridge, Hills Rd, Cambridge CB2 0AH; National Research Council, Institute of Genetics and Biophysics, Naples, Italy

## Abstract

Purging through inbreeding occurs when consanguineous marriages increases the rate at which deleterious alleles are present in a homozygous state. In this study we carried out low-read depth (4-10x) whole-genome sequencing in 568 individuals from three Italian founder populations, and compared it to data from other Italian and European populations from the 1000 Genomes Project. We show extended consanguinity and depletion of homozygous genotypes at potentially detrimental sites in the founder populations compared to outbred populations. However these patterns are not compatible with the hypothesis of consanguinity driving the purging of highly deleterious mutations according to simulations. Therefore we conclude that genetic drift and the founder effect should be responsible for the observed purging of deleterious variants.

## Introduction

Population genetics theory and empirical evidence from model organisms predict that deleterious variants are purged by selection at a rate proportional to their fitness effect and the effective population size^1^. However, evidence for similar genetic load in populations that underwent bottlenecks and constant size populations shows that selection is equally effective regardless of demography, and therefore a reduction in population size has no effect on the population's genetic load^2,3,4^. Nevertheless, other demographic factors can affect the efficacy of selection. Inbreeding depression is the reduction in average population fitness due to higher level of homozygosis than expected under Hardy-Weinberg equilibrium. Increased homozygosity can lower fitness either by reducing the heterozygous genotypes at loci with heterozygous advantage, or by increasing the rate of homozygosity for partially recessive detrimental mutations^5^. This last case exposes deleterious mutations to selection, driving their purging from the population increasing the effectiveness of selection.^6,7,8^ In cases of systematic inbreeding due to non-random mating (as is the case of small consanguineous populations), purging can work even for mildly deleterious alleles. However, purging is effective only for lethal or semilethal alleles in the context of as opposed to cases of panmictic inbreeding (i.e. inbreeding due to finite size of the populations) and drift^9^.

Consanguineous marriages result in substantial deviation from Hardy-Weinberg equilibrium due to increased autozygosity, which can cause deleterious recessive variants to be present in a homozygous state. While autozygosity is associated with higher rate of disorders (congenital, late onset, infertility, miscarriage, infant mortality and morbidity) in consanguineous populations^10–14^, it also provides an opportunity for selection to act by purging exposed deleterious phenotypes, eventually reducing the burden of homozygous deleterious genotypes^8^. Purging causes less fit individuals to be eliminated, ultimately increasing the population's fitness. Indeed, in some species inbreeding caused by a bottleneck has been shown to have beneficial effects, for instance by promoting the invasiveness of some species of ladybirds^15^ or removing highly deleterious variants in mountain gorillas^16^. In humans, consanguineous couples experience higher fertility than non-consanguineous couples ^12,17,18^, but this has been associated with causes other than purging of deleterious variants, such as social factors (e.g. younger maternal age at first birth), reproductive compensation^12^, and the negative effects of outbreeding, including the break up of coadapted gene complexes, and the increase of maternal-foetal incompatibility^18^. Little is known about how inbreeding might play a role in the improved fecundity of consanguineous couples. In this study we investigated whether autozygosis has had a role in the purging of deleterious mutations in three Italian founder populations with different demographic histories^19–21^ by comparing them with outbred Italian and European populations. We show evidence for a depletion of homozygous genotypes at putatively detrimental sites in all the three populations. The presence of this effect in multiple founder populations with high rates of consanguinity suggested that inbreeding might have had a role in accelerating purging of deleterious mutations. However, according to simulated results consanguinity could not be responsible for the observed purging, therefore we conclude that genetic drift and the founder effect have driven the observed purging of deleterious variants from these isolated human populations.

## Results

### Whole genome sequence of isolates identified 21M variable sites

We carried out low-read depth whole-genome sequencing (average 6x, Table S1) in 568 unselected individuals from three populations from the Italian network of genetic isolates (INGI), namely Val Borbera (VBI), Carlantino (CAR) and Friuli Venezia Giulia (FVG). Of these, FVG is further subdivided in 4 different villages, namely Erto (FVG-E), Illegio (FVG-I), Resia (FVG-R) and Sauris (FVG-S) (Figure 1a and Table S2). After stringent quality control (Methods), we identified a total of 21,244,190 variable sites. We further compared genetic variability of the isolates with populations representative of the general Italian and European population, namely 98 Tuscans (TSI) and 85 Utah Residents with Northern and Western European ancestry (CEU) from the 1000 Genomes Project (TGP)Phase 1 data set^22,23^ (Figure 1a). The isolate call sets were generated using the same variant calling and QC pipelines employed by the TGP project, so the two variant datasets were merged to generate a final set of 46,281,641 variable sites including sites variable both in isolates and TGP. To verify the validity of this approach, we estimated concordance of genotypes at 298k variable sites on chromosome 22 within the merged call set with those obtained by simultaneously calling isolates and the TGP. We found a median non-reference genotype discordance of 1.55% and 3.8% for SNPs and INDELs, respectively (Table S3), suggesting that merging the two call sets is a viable approach.

**Figure 1.**
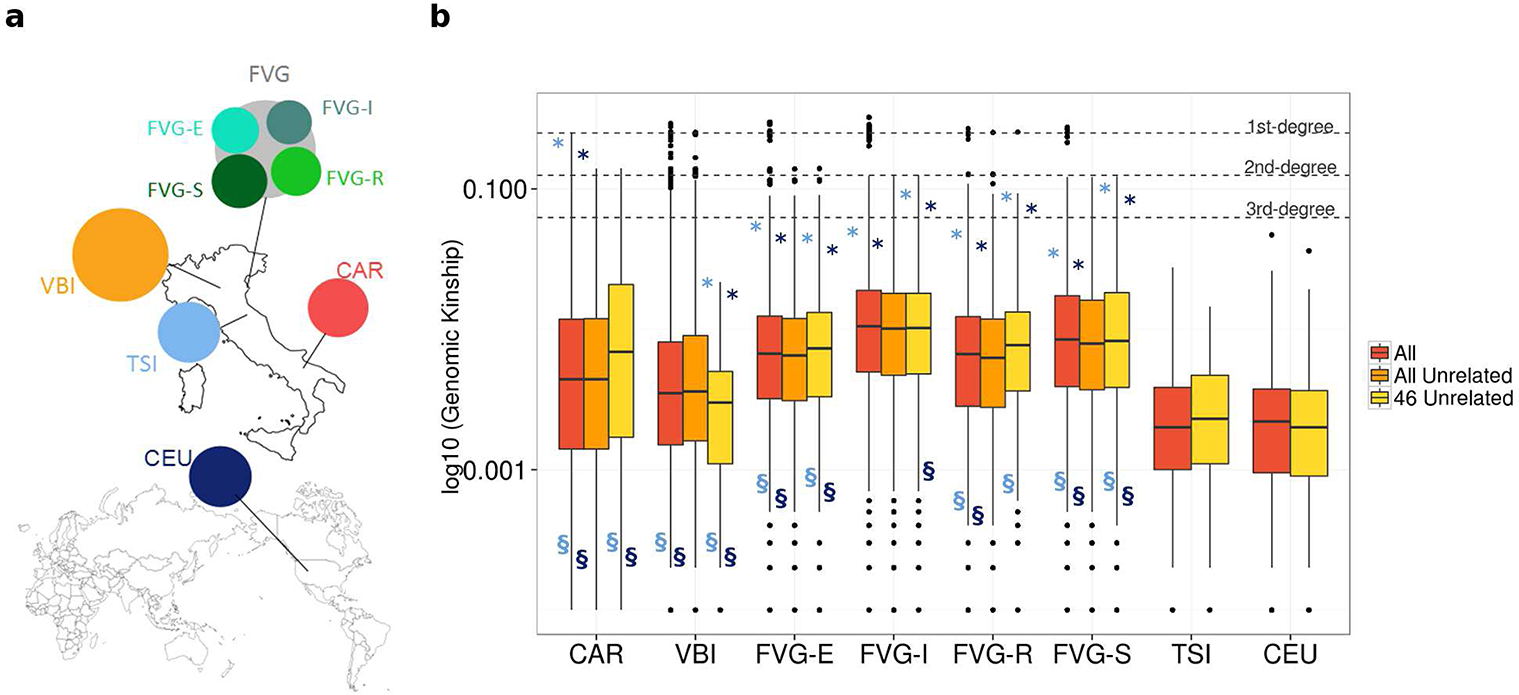
Geographical origin of the population and evidence for systematic consanguinity in the isolated populations. **(a)** Geographic location of the populations in this study. Circles are proportional to sample sizes. VBI = Val Borbera, CAR = Carlantino, FVG = Friuli Venezia Giulia, FVG-E = Erto, FVG-I = Illegio, FVG-R = Resia and FVG-S = Sauris. The figure was drawn using GNU Image Manipulation Program using maps available in the public domain. (b) Boxplots of genome-based kinship distributions. In each population we measured genomic kinship for all possible pairs derived from: (i) all individuals (“All”); (ii) all individuals after removing one individual per pair in pairs related to 1st degree (“All Unrelated”); (iii) 46 random individuals after checking that they were not related to 1st degree in isolates. The category “All Unrelated” is empty for CEU and TSI, as there were no pairs related to 1st degree in these populations. Grey dashed lines represent reference values for 1st-, 2nd- and 3rd-degree relationships. Dark and light blue * and § indicate significance in comparisons with CEU and TSI, respectively (p-values in Table S5). Individuals in isolates tend to be more consanguineous than reference populations. In FVG-R and VBI relatedness to 1st degree persists even after removing 1st degree relatives according to pedigree data. The explanation is that ancestors of some 2nd and 3rd degree relatives were in turn the results of consanguineous relationship, consistent with long term consanguinity in isolates.

We investigated genetic relationship among individuals of the three isolates, TSI and CEU using a random set of 281k variable chromosome 22 sites spanning all frequency ranges. A previous exploration of VBI and FVG was limited to low-resolution SNP array data which is not informative for rare, private and novel genetic variation^19,20^ while the genetic structure of CAR has not been described before. In a principal component analysis, two FVG villages (FVG-R and FVG-S) appeared to be distinct from the other populations along the first three components. Further, VBI and CAR separated from CEU and TSI on the fourth component (Figure S1). In admixture analyses, FVG-R and FVG-S separated from other populations already at K=4 clusters, although the most likely number of clusters was 10 (Figure S2). These results confirm earlier results based on SNP array data from the entire FVG sample^20^, and are thus not related to the sampling strategy used in this study. Because of this sub-structuring, when relevant FVG villages were considered separately for subsequent analyses. CAR, described here for the first time, clusters with VBI and other reference populations (TSI and CEU) along the first two components and with VBI along the 3^rd^ and 4^th^ component (Figures S1 and S2).

Using a different data set of variants obtained by jointly calling the TGP and isolated populations cohorts that do not include CAR, we evaluated FST distance between pairs of populations (Table S4). As shown in previous publications^20^ the between-populations pairwise Fst values of the FVG villages were extremely high. The overall mean between pairs of villages is 2%, which is substantially higher than the Fst values between the villages and the other populations, which range from 1-1.5%. Other pairwise Fst were below .5%.

### Long term isolation and diffused homozygosity in isolated populations

FVG^20^ and VBI^21,19^ are located in two mountain regions of North Italy that were in the past poorly connected to neighbouring areas. In addition to geographical isolation, FVG villages experienced linguistic isolation due to their use of very different dialects. In the 14th century people moving from the area of Naples migrated some 300 km to found the CAR population, which experienced isolation due to differences in dialect with surrounding communities and poor connection to its parent population. The historical isolation experienced by isolates causes systematic inbreeding, which is expected to result in greater genetic homogeneity and homozygosity as compared to reference population. We investigated this hypothesis using several approaches.

First, we calculated relatedness between pairs of individuals using two independent measures, namely pedigree-based and genome-based kinship. Pedigree based kinship represents the expected fraction of the genome shared that is identical by descent (IBD) and was calculated from available genealogical records dating back to the 17th century for VBI ^21,24^, records collected during sampling of DNA for FVG, and from private genealogical collections available for CAR. Genomic kinship is the fraction of the genome which is shared identical by descent and was calculated using genetic data. Pedigree- and genome-based kinship values were highly correlated (Figure S3), however in FVG villages the resolution of pedigree-based kinship was poor because genealogies are incomplete, therefore we focused on genomic based-kinship.

Genome-based kinship values are significantly higher in isolates than reference populations and average kinship in isolates is always higher than reference populations (Figure 1b; Table S5; two-sample Kolmogorov-Smirnov and Mann-Whitney test p-value in Table S6). The kinship distributions were highly skewed towards low values in all cases, however maximum values were in the range of 1^st^ degree relationships for isolates, compared to approximately 3^rd^ degree in reference populations (Figure 1b, “All”). When removing one individual per pair from pairs related to the 1^st^ degree according to pedigree data, we still observed genomic-kinship values in the range of 1^st^ degree in FVG-R and in VBI (Figure 1b, “All Unrelated”). We confirm that the observed differences were not due to different sample sizes, by re-calculating genomic kinship for a subset of 46 random individuals per population (Figure 1b, “46 Unrelated”). Overall, these results not only show that individuals from isolates tend to be more consanguineous than reference populations, but also suggest that consanguinity has systematically occurred in these populations in historical times, and ancestors of 2^nd^ and 3^rd^ degree relatives were in turn the result of consanguineous relationships. Because of this extended consanguinity, we excluded from subsequent analyses one randomly-selected individual from pairs with 1^st^ degree relationships (Table S1)

Having assessed diffused consanguinity in the isolates, we next quantified genome segments which were IBD among pairs of 46 random unrelated individuals per population. In Figure 2a we report the distributions of total extension of IBD per genome (i.e. the sum of single segments). While there was no significant difference between isolates and CEU or TSI in the total IBD distributions (two-sample Kolmogorov-Smirnov test p-value >0.01, Table S7) FVG villages displayed higher total IBD sharing on average (Figure S4 and Table S7). These differences were more evident when contrasting cumulative distributions of total IBD (Figure 2c and Table S7): in FVG villages 95% of the individuals shared 337–461(+/-85–123)Mb compared to 122–157(+/-8–45)Mb in other isolates and reference populations. Considering an accessible genome size of 2.5Gb^23^, this would indicate that individuals in FVG villages share between 13% and 18% of their genomes, *versus* 5–6% in other populations, corroborating evidence for extended consanguinity in the isolates.

**Figure 2.**
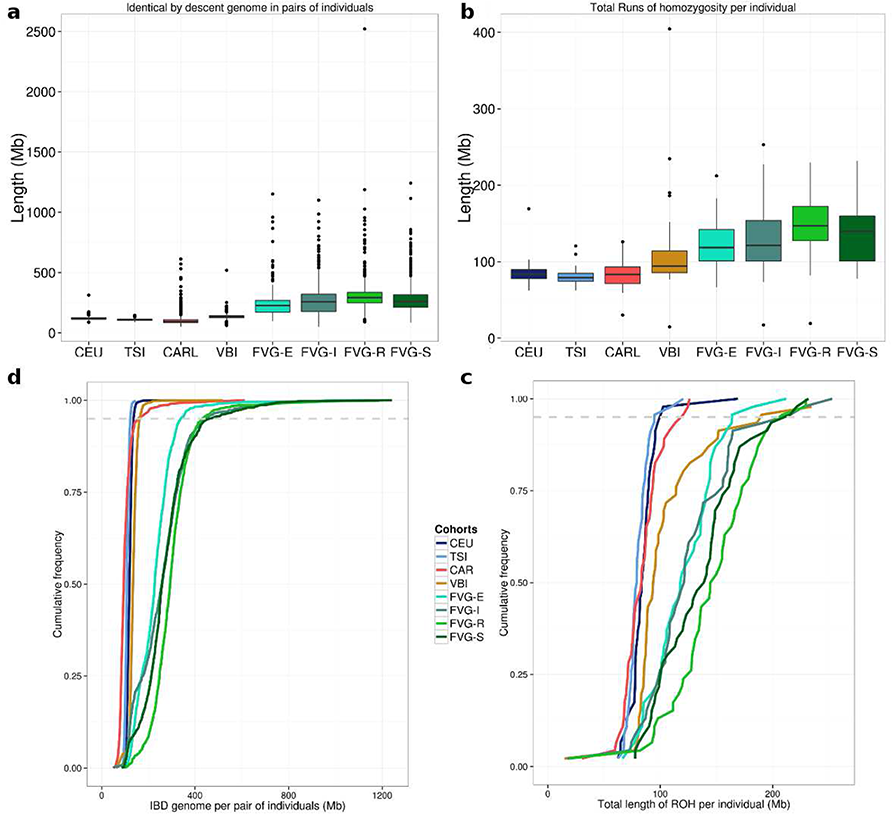
Higher genome sharing between individuals and within individual in isolates compared to outbred reference populations. Summary statistics **(a and b)** and cumulative frequency **(c and d)** of the total length of segments identical by descent (IBD) per pairs of individuals and total length of runs of homozygosity (ROH) per single individual. In both (c) and (d) the dashed line indicates the 95% percentile. Considering an accessible genome size of 2.5Gb, individuals in FVG villages share between 13% and 18% of their genomes, versus 5–6% in other populations (c) and 5–8% of the genome in is in homozygosity in isolates (except CAR) compared to 3–4% in CEU, TSI (d).

Finally, we investigated allele sharing within an individual by identifying genomic segments with contiguous genotypes in homozygosity (Runs Of Homozygosity, ROH) in each individual. We report in Figure 2b the distributions of the total length of ROH (i.e. the sum of single segments) per individual. With the exception of CAR, ROH extension was significantly higher in isolates compared to CEU and TSI (two-sample Kolmogorov-Smirnov test p-values in Table S7), suggesting increased homozygosity in isolates. This is also evident when comparing cumulative distributions of total ROH in 46 randomly selected individuals per population (Figure 2d, Table S7): 95% of individuals in FVG villages and VBI had between 163–209(+/-30–43)Mb of genome in homozygosis compared to less than 100(+/-10–15)Mb in the outbred reference populations. These figures suggest 5–8% of the genome is in homozygosity in isolates (except CAR) while only 3–4% is in CEU and TSI.

When ROHs segments were classified according to their length^25^, we observed a greater prevalence of medium and long ROHs in isolates (and especially FVG-R and FVG-S) than in CEU and TSI (Figure S5). This condition is consistent with the presence of extended consanguinity or very recent admixture; both cases apply to the isolates under study here, for which isolation (either geographical or linguistic) was a past condition that is no longer met.

### Isolates experienced genetic drift and a recent decrease of the effective population size

Isolated populations are expected to be more greatly subjected to the effect of genetic drift compared to outbred populations, therefore we investigated to what extent genetic drift acted on the populations in this study. We used a method based on inference from genomic segments that are shared IBD which enables the exploration of very recent fluctuations in effective population size^26^. Low coverage and computational phasing are likely to affect the accuracy of IBD detection methods, occasionally breaking down long IBD segments into shorter ones and thus potentially creating the artefactual effect of a population expansion. Despite this potential confounding, our analysis displayed clear patterns of overall lower effective population size estimates in isolates compared to reference populations (Figure 3). In the last fifty generations (approximately last 1500 years) we observed a population size contraction in FVG villages compared to CEU and TSI, which appear unaffected by the recent trends towards expansion of other European populations^27,28^. Such abundant sharing of long haplotypes co-inherited from recent common ancestors during a recent population contraction provides further evidence for genetic drift that extended until the very recent centuries in these isolated groups. CAR and VBI have a less pronounced trend of contraction compared to FVG, however they still have smaller effective population sizes compared to TSI and CEU. We therefore conclude that isolates experienced more genetic drift compared to reference populations.

**Figure 3.**
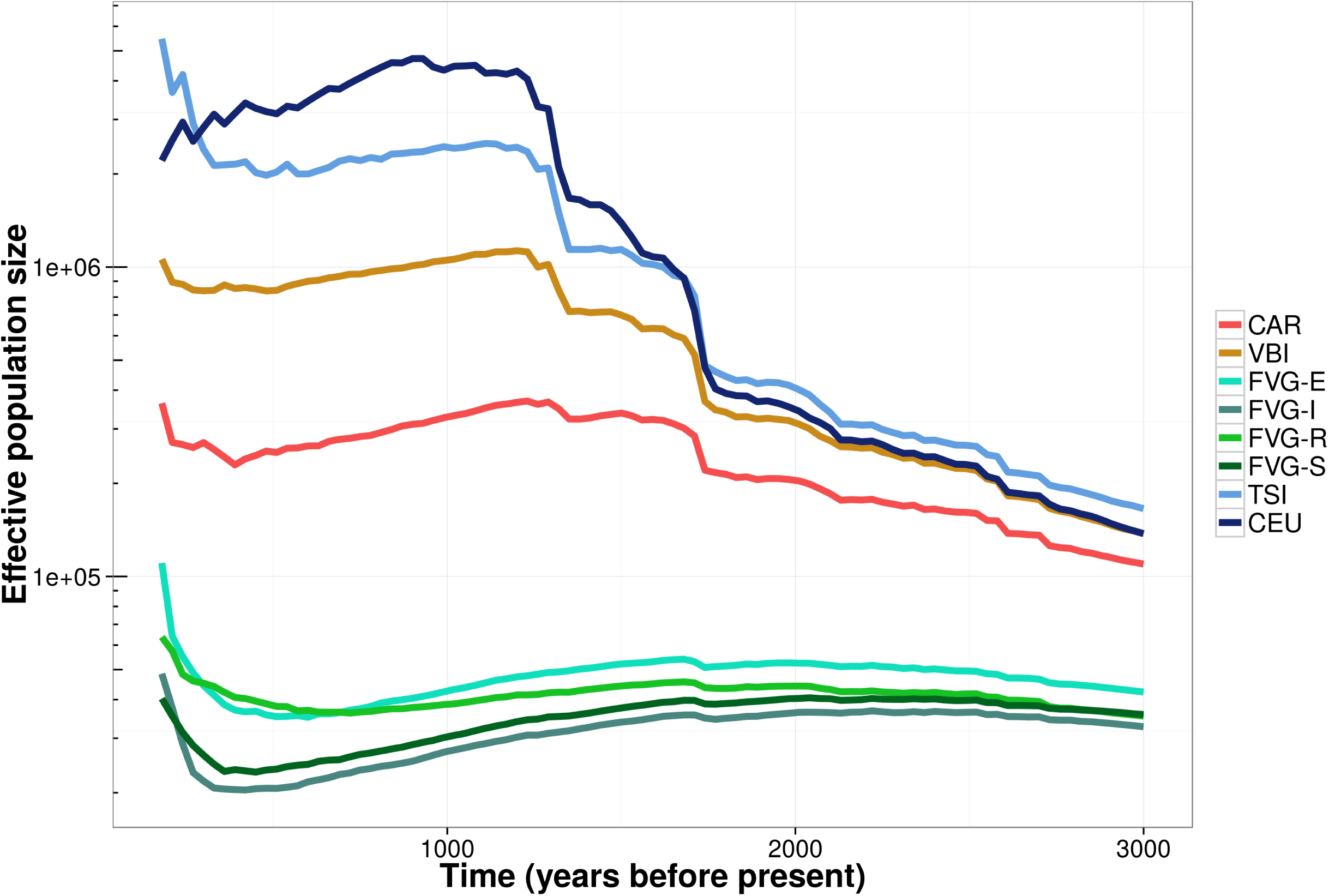
Isolates experienced more genetic drift compared to reference populations. Trends of the effective population size in the last 3000 years inferred from identical by descent genome of individuals within populations. In the last fifty generations (last ~1500 years) there was a population size contraction in FVG villages compared to CEU and TSI, which appear unaffected by the very recent trends towards expansion of other European populations. VBI and CAR instead follow the trend of CEU and TSI but with substantial smaller effective population size.

### Depletion of putatively detrimental sites in isolates

To test the hypothesis that autozygosity contributes to the purging of deleterious mutations through exposure of detrimental alleles to purifying selection, we determined the incidence of homozygous detrimental genotypes and measured the efficiency of purifying selection in isolates compared to reference populations.

To compare allele frequencies we used a different data set of variants obtained by jointly calling the TGP and isolated populations cohorts and do not include CAR. Starting from this data set we selected 322 individuals (46 randomly selected unrelated individuals per population) for which we identified 12,344,818 variable sites. Of these, 10,917,001 sites had information on the ancestral state of the alleles and were considered for further analyses. After determining derived allele frequencies (DAF) per each population we evaluated incidence of derived allele causing missense (356,155 sites), synonymous (229,529 sites), or loss of function (LoF, 4,286 sites)^29^ consequences. Alleles determining missense (and some time synonymous^30^) consequences introduce changes to the coded protein that might alter the protein structure, whereas LoF alleles completely disrupt the protein function. We used a million sites annotated as intergenic as a proxy for neutrality in comparisons.

With few exception, we observed significantly fewer derived alleles counts per individual in FVG and VBI compared to reference populations at synonymous and intergenic sites, whereas derived allele counts at missense and LoF sites was overall comparable (Figure 4a, Wilcoxon rank sum test p-values in Table S8). We then evaluated how often putatively detrimental alleles are found in homozygosis by counting the number of homozygous derived genotypes per individuals. Isolates had significantly higher rates of homozygous derived genotypes at neutral and synonymous sites (Figure 4b, Table S8) compared to reference populations, in accordance with the general homozygosity trend reported in a previous section. However, with one exception (FVG-R), we found no significant differences in the rates of homozygosity at missense and LoF deleterious sites between the reference and isolate populations. This is indicative of an overall depletion of homozygous genotypes at deleterious sites in the isolates, which, along with the comparatively high rates of homozygosis at neutral alleles, supports the hypothesis the hypothesis that deleterious alleles have been purged through inbreeding in the isolates

**Figure 4.**
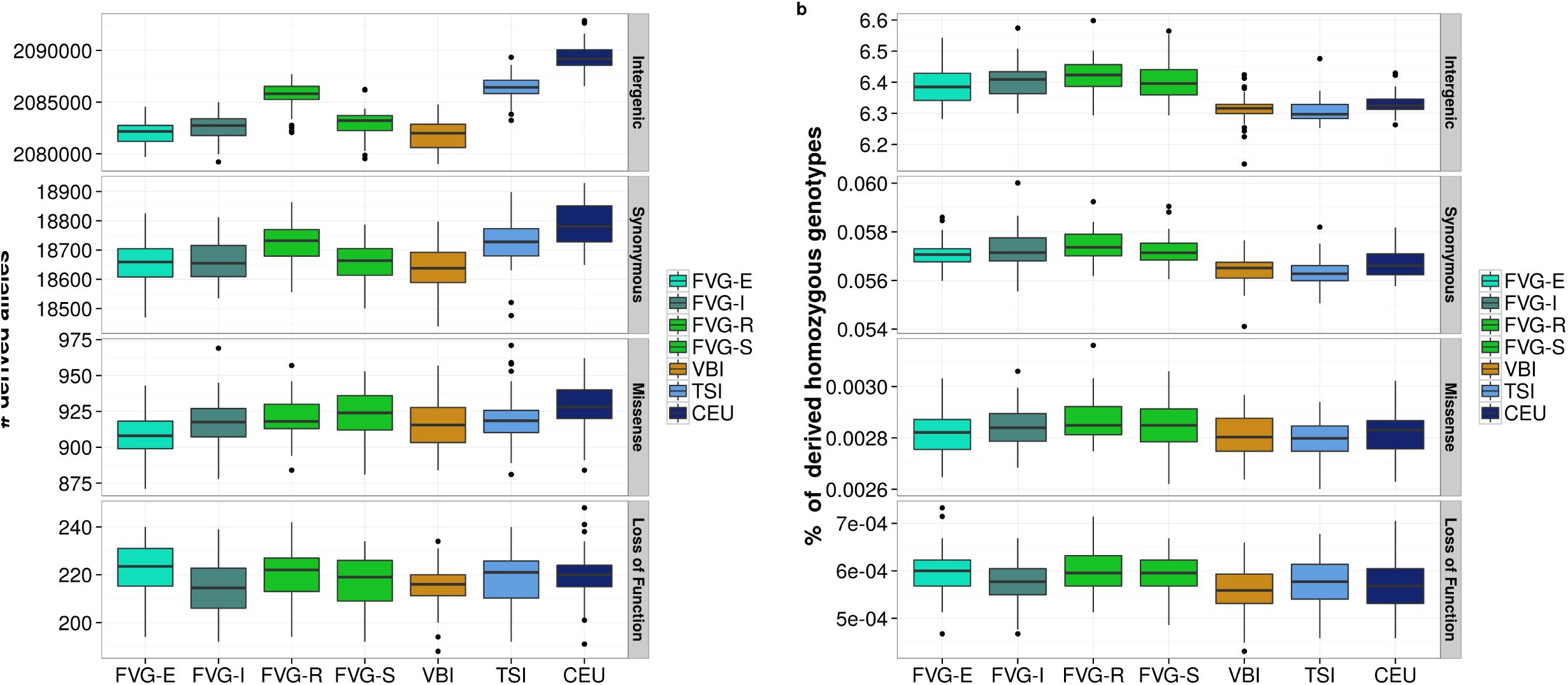
Depletion in the isolates of homozygous genotypes at missense and loss of function (LoF) variants. **(a)** Counts of derived allele per individual at variable sites with intergenic, synonymous, missense and LoF consequences. Statistically significant comparisons (p-value<0.001, Table S7) of isolates with CEU and TSI are marked by dark and light blue asterisks, respectively. In the isolates individuals have less derived alleles compared to reference populations at intergenic and synonymous sites. (b) Fraction of genotypes homozygous for the derived allele at the same functional categories as in (a). At intergenic and synonymous sites isolates have significantly higher rates of homozygosity compared to reference populations (Table S7) whereas at missense and LoF sites rates are comparable, suggesting a depletion of homozygous deleterious genotypes in isolates.

We therefore used a recently proposed metric (R_X/Y_)^2,4^ to assess whether selection has been more effective in isolates compared to reference populations. The metric R_X/Y_ contrasts frequencies of alleles between pairs of populations and measures their accumulation in one population compared to the other: a value of 1 indicates that the two populations have comparable number of mutations per individual, whereas values lower than 1 indicate a depletion in population X and values higher than 1 indicate a depletion in population Y. We compared R_X/Y_ at intergenic, synonymous, missense, and LoF variants in isolates and reference populations using derived allele counts of 46 randomly selected individuals per population.

Overall, R_X/TSI_ values for all functional categories were closer to one compared to R_X/CEU_ ones, reflecting the greater similarity of isolates to TSI than to CEU (Figure 5). With the exception of VBI and FVG-I, in all comparisons involving intergenic and LoF sites R_X/Y_ was not significantly different from one, however for LoF R_X/Y_ values were less accurate, because of the low number of sites considered. In most cases, R_X/Y_ at synonymous and intergenic sites was similar and a trend of diminishing R_X/Y_ is visible among intergenic, synonymous and missense sites. From a populations perspective, for VBI both R_X/CEU_ and R_X/TSI_ were always significantly less than one at all functional categories, whereas for FVG villages patterns are less simple. Overall, besides details of single categories and populations, we observed many cases in which R_X/Y_ was significantly lower than one in isolates especially at sites with deleterious functional consequences (i.e. missense) suggesting that isolates were depleted of putatively detrimental alleles. While for intergenic sites R_X/Y_ is almost always significantly equal to 1, at other categories it is often significantly <1 suggesting more efficient selection in isolates compared to reference population. These patterns could possibly arise due to the founder effect, the genetic drift and the purging effect of inbreeding alongside strong selection coefficients for highly disruptive mutations.

**Figure 5.**
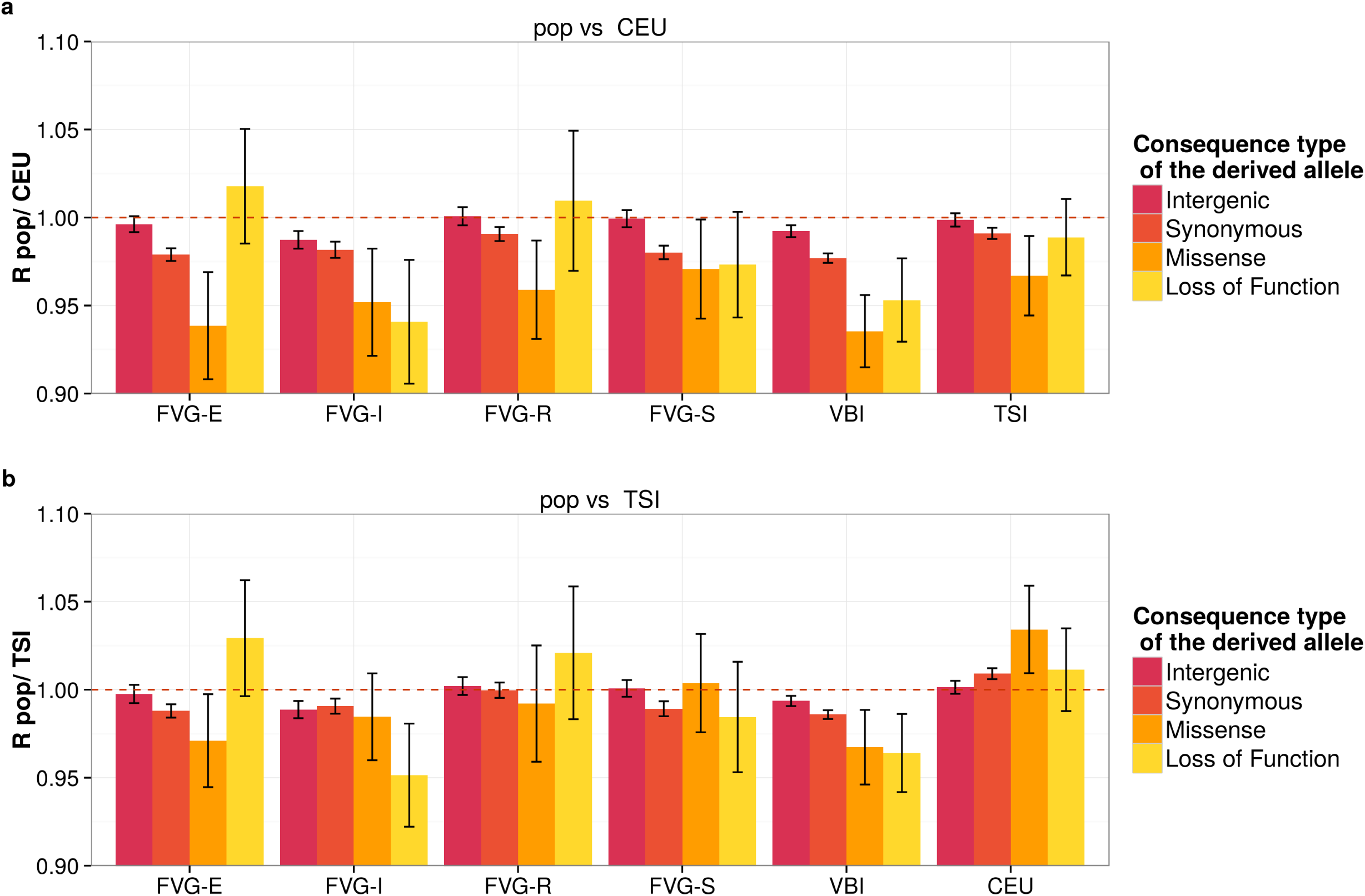
Isolates have less missense alleles compared to reference populations. Pairwise comparison of isolates with CEU **(a)** and TSI **(b)** at missense, synonymous and LoF sites. RX/Y is a measure of the accumulation of mutations in population X respect to population Y; values lower than 1 indicate a depletion in population X and viceversa. Bars represent variance obtained from block jackknifes. While for intergenic sites RX/Y is almost always significantly equal to 1, at other categories it is often significantly <1 suggesting more efficient selection in isolates compared to reference population. Possible causes are the founder effect, the genetic drift and the purging effect of inbreeding alongside strong selection coefficients for highly disruptive mutations. At LoF sites RX/Y are not significantly different from one, however for LoF RX/Y values were less accurate, because of the low number of sites considered compared to other categories.

### Simulations support the observed depletion of highly deleterious homozygous genotypes but fail to see any effect of inbreeding

We evaluated expectations to observe purging in simulated data under a range of demographic scenarios, natural selection intensities and models. A population isolate was simulated to arise from an European-like population at 160, 60 or 20 generations ago (generation time 25 years), and to evolve to present time under five demographic models (continued bottleneck, serial founder effect, bottleneck plus linear expansion, bottleneck plus instantaneous expansion and bottleneck plus linear reduction; Figure S6) and three different values of average kinship (0, 0.002 and 0.01). We forward-in-time simulated genomic data by mimicking features of exomes spanning 115 kbp (Methods). Mutations were assumed to be deleterious ¾ of the time in exons and ½ in UTRs under six selection coefficients *(s* = -0.2, -0.1, -0.05, -0.01, -0.005 and -0.001). Each scenario was replicated 500 times. We found purging to be more effective in the bottleneck plus expansion (Figure S7) and thus for this model we further explored a range of bottleneck intensities (effective population size, Ne, during the bottleneck = 15, 25 and 50 and bottleneck duration 1 to 25 generations in steps of 5, Table S9). Demographic parameter choice was a compromise between computational feasibility and similarity to the demography of the isolates under study, its main purpose being to explore properties of purging rather than to perfectly fit isolate demography.

Purging was calculated in a sample of 50 individuals as the fold change between present and past (at founding event) number of recessive deleterious homozygous genotypes per individual. A negative fold change indicates a reduction of deleterious homozygous genotypes, as is expected in the case of purging. Results in Figure 6 (also see Figure S8) show that: (i) under neutrality (*s* = 0) no purging is seen in all three models; (ii) for mildly deleterious mutations (*s* = 0.01) purging is almost complete under dominance, moderate for additive models but not observed under recessive models; and (iii) for highly deleterious mutations (*s* = 0.1) purging of homozygous recessive deleterious genotypes is also seen under recessive models.

**Figure 6.**
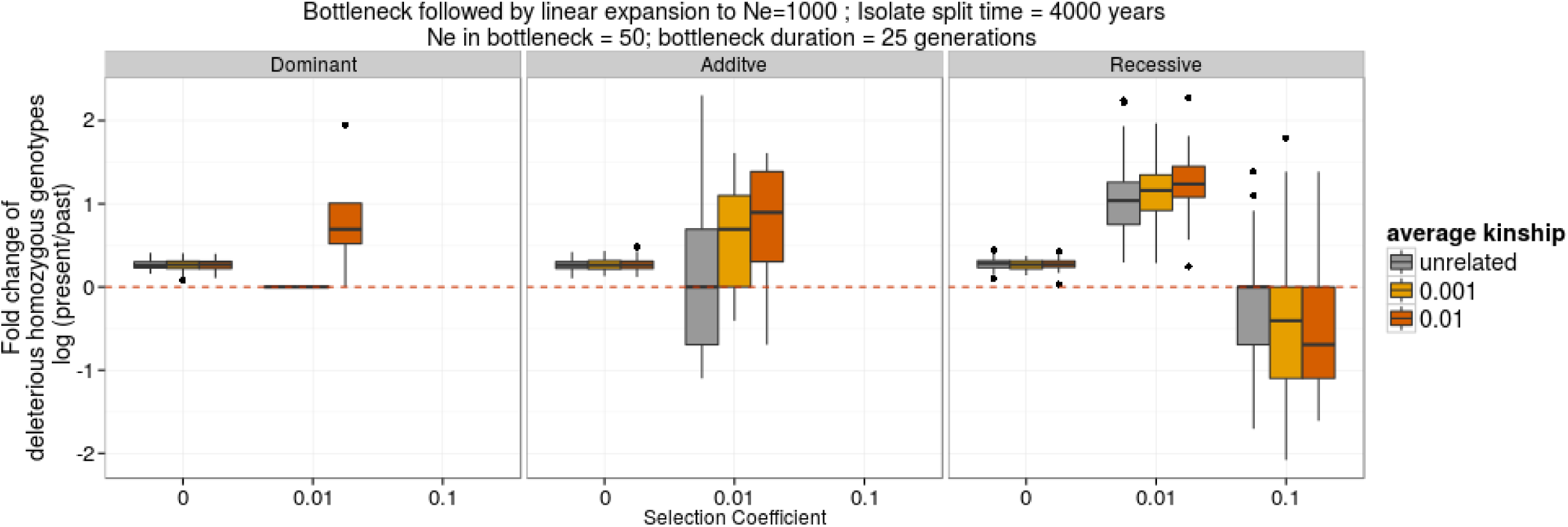
No contribution of the inbreeding to the depletion of deleterious mutations according to simulated results. Purging is measured as the fold change between present and past (at founding event) average number of recessive deleterious homozygous genotypes per individual. A negative fold change is expected in case of purging. Boxplots summarize results of 100 replicates, each replicate being the average over 50 individuals. Dominant, additive and recessive models are shown for neutral (selection coefficient, s=0), mildly deleterious (s=0.01) and highly deleterious (s=0.1) variants in three populations with different average kinship. Ne=effective population size.

Overall, these simulations suggest that the selection coefficient and dominance model have a major effect on purging and that purging is possible for highly deleterious recessive mutations. While it is difficult to make assumptions on the inheritance model of missense and synonymous variants, a recessive model and a strong selection coefficient are highly plausible for LoF^29^. We therefore conclude that simulations support the observed depletion of highly deleterious homozygous genotypes observed at LoF sites. Nevertheless, our set of simulations does not explain the role of consanguinity as the effect of kinship is negligible (Figure 6). This may be due to the imperfect fit of the demographic parameters used for simulations to the isolates studied here. A more comprehensive analysis will be required to fully understand this aspect.

## Discussion

Inbreeding consists of mating between closely related individuals. Chromosome pairs of inbred offspring share segments that are identical by descent more often than expected by chance, and consequently large portions of the genome in inbred individuals are in homozygosis. Inbred offspring undergo inbreeding depression, the phenomenon by which the overall fitness of the population is reduced due to higher levels of homozygosity than Hardy-Weinberg expectations, including the rise of homozygous detrimental mutations. However, inbreeding depression cannot be maintained permanently, as eventually the genetic load caused by the rise of the frequency of detrimental mutations is offset by an increased efficiency of selection as inbreeding drives higher rates of homozygosis at recessive deleterious alleles, increasing the chance that these alleles manifest a phenotype that may be selected upon in a process known as genetic purging. Therefore, in the long term inbreeding can have positive effects on fitness. While this was demonstrated to be the case for other species (e.g.^15,16^), it has never been demonstrated for humans^2^.

We decided to assess if genetic purging has occurred in three Italian isolated populations which are likely to feature high rates of inbreeding. By performing the analysis of whole genome data we first demonstrated the occurrence of (i) diffused consanguinity, (ii) extended regions of autozygosity and (iii) small effective population sizes until recent times, showing that indeed the three populations are genetic isolates and the genomes of their members are more homogeneous than those of representatives of outbred European and Italian populations. Secondly, we compared allele frequencies between isolates and reference populations at putatively detrimental and neutral sites and showed a depletion of homozygous genotypes of putatively deleterious mutations in the isolates. Furthermore, according to a statistic that measures the efficacy of selection at removing deleterious genotypes, population isolates have fewer deleterious allele at detrimental sites than neutral ones.

Following this evidence we then used simulations to understand if the depletion of homozygous deleterious genotypes could have been the result of purging using simulations. In accordance with other published studies^4^, our simulations showed that a reduction of homozygous deleterious genotypes is indeed possible after a bottleneck assuming a recessive model and strong selective pressure. However, we did not observe a change in the efficacy of selection when varying the inbreeding coefficient of the simulated population. Therefore we concluded that under the considered simulation scenario, the observed depletion of deleterious homozygous genotypes might be a general effect of the demography compatible with the general loss of rare alleles due to bottlenecks and genetic drift.

Consanguineous populations represent a model for the effects of systematic inbreeding. Consanguinity can naturally occur for several reasons, including geographical, linguistic and religious isolation. Recently there has been a great interest in exploring the genetic load in populations with different demographic history. Consanguineous populations in this study complement the range of demographic scenarios and here we demonstrate that they provide an excellent model study to understand the interplay between selection, genetic drift and homozygosis in determining the genetic load of human populations.

## Methods

### Samples individuals and data generation

Low coverage whole genome sequence was generated in for 568 individuals belonging to the INGI network. Individuals are from Italian populations form North-West (Val Borbera^21^, 225 samples), North-East (Friuli Venezia Giulia^20^, 250 samples) and South-West (Carlantino^20^, 93 samples). Sequencing was carried out using Illumina technology (Genome Analyzer and HiSeq 2000) at the Wellcome Trust Sanger Institute and Beijing Genomics Institute (54 samples from FVG cohort). Data coverage was 4-10X (Table S2).

The study was reviewed and approved by the following Ethical Committees: Ethical committee of the San Raffaele Hospital and of the Piemonte Region (VBI), Ethics approval was obtained from the Ethics Committee of the Burlo Garofolo children's hospital in Trieste (FVG), the local administration of Carlantino, the Health Service of Foggia Province, Italy, and ethical committee of the IRCCS Burlo-Garofolo of Trieste (CAR). Written informed consent was obtained from every participant to the study and all methods were carried out in accordance with the approved guidelines (Declaration of Helsinki of the World Medical Association).

### Data processing, variant calling

Genotype calls were produced for each population separately using the following pipeline. Samtools mpileup (v. 0.1.19)^31^ was used for multisample genotype calling (parameter set: -EDVSp -C50 -m3 -F0.2 -d 40). Variant Quality Score Recalibrator (VQSR) filtering was applied to the raw call data with GATK v.2.5^32^ through VariantRecalibrator module separately for SNVs and INDELs. The filter creates a Gaussian Mixture Model by looking at annotations values over a high quality subset of the input call set and then evaluate all input variants. For SNVs we used the following parameters: I) Annotations: QD, DP, FS, HaplotypeScore, MQRankSum, ReadPosRankSum, InbreedingCoeff II) Training set: HapMap 3.3, Omni 2.5M chip, 1000 Genomes Phase I; III) Truth set: HapMap 3.3, Omni 2.5M chip; IV) Known set: dbSNP build 138. For INDELs we selected: I) Annotations:

DP,FS,ReadPosRankSum,MQRankSum; II) Training set: Mills-Devine, 1000 Genomes Phase I, DbSnp v138; III) Truth set: Mills-Devine; IV) Known set: Mills-Devine, dbSNP build 138. For each population the lowest VQSLOD threshold has been chosen according to the output produced by VariantRecalibrator to select the best cut off in terms of specificity and sensitivity of the trained model. For SNPs the minimum VQSLOD values selected are 3.016 (97.3% truth sensitivity threshold), 2.8309 (97.82% truth sensitivity threshold), 1.4512 (98.5% truth sensitivity threshold) for FVG, VBI and CARL cohort respectively. Since INDELs calling and alignment is still more prone to error we used a conservative approach, selecting a sensitivity threshold of 90% for each population. The filter has been applied using GATK's Apply Recalibration module.

To ameliorate the low quality of raw genotyping results based on the low coverage sequencing data we performed several genotype refinement steps on the filtered call set: 1)we used BEAGLEv4.r1230^33^ to assign posterior probabilities to all remaining genotypes. 2) SHAPEITv2^34^ has been used to phase all genotypes call and 3) IMPUTEv2^35,36^ to perform internal imputation in order to correct genotyping errors. Information about Ancestral Allele and allele frequencies from TGP populations have been retrieved from dbSNP v.138^37^. The Variant Effect Predictor v.74^38^ provided all consequence annotation as well as Polyphen,Sift and GERP informations.

We merged genotype call set of the INGI populations with low coverage data available from Phase1 of the 1000 Genomes Project^23^ in a union using set using bcftools merge^31^. To check if this approach would introduce bias in downstream analyses, we evaluated genotype discordance on chromosome 22 between the union set and a genotype call set in which variants were called simultaneously in all INGI, CEU and TSI populations using the same pipeline described above. Discordance was calculated by comparing the genotypes of the same sample in the two sets and counting cases in which alleles were called differently, e.g. the Ref/Ref_discordance was the proportion of cases where a Ref/Ref genotypes in one set was wrongly called as a Ref/Alt or Alt/Alt in the other set. We details these definitions in the Table S3 caption.

### Joint calls from the data set of the Haplotype Reference Consortium

The Haplotype Reference Consortium (HRC.r1) callset was created by merging the data from 20 separate large whole-genome sequencing projects (see http://www.haplotype-reference-consortium.org for details of these cohorts including numbers and average sequencing coverage). The initial set of sites was selected by combining the individual phased SNP callsets from these cohorts. Non-biallelic sites were split into biallelic sites and sites with minor allele count (MAC) >= 5 were selected. Genotype likelihoods (GLs) were extracted from the original alignments (BAM files) using samtools mpileup/bcftools call^31^. Having generated GLs at these sites, further site filtering was performed by removing a set of sites that could be putatively called within cohorts from the likelihoods, but have been filtered out consistently across the initial callsets. The HRC.r1 sites were further filtered sites for which any of the following statements were true: (i) Per-cohort Hardy-Weinberg Equilibrium (HWE) p-value < 10^10^ and cohort is not 1000 Genomes; (ii) Overall inbreeding coefficient < -0.1; (iii) Minor allele frequency (MAF) > 0.1, site called in less than three cohorts, and site not called in 1000 Genomes. Additionally sites found on a selection of common SNP genotyping arrays were reintroduced if they had been filtered by any of these steps. The final site list contains 39,235,157 autosomal sites. In Table S10 we report fractions of sites with p(HWE) <1.0e-4 per population. Allele frequency spectrum for the minor allele is shown in Figure S10.

From the initial set of samples collected, some were removed due to low call rate based on putative calls from bcftools based on GLs. A further set of samples were removed to eliminate duplicate samples that had been sequenced across multiple studies, resulting in 32,488 samples in the final set. A new extension of the SNPtools algorithm^39^ which takes advantage of pre-existing haplotypes was used to call haplotypes from the GLs. We merged these haplotypes with the pre-existing haplotypes by setting the genotypes of any sites which were only found in the pre-existing haplotypes to be homozygous major allele in the new haplotypes. A development version of SHAPEIT was then used to refine the phasing of the called haplotypes.

### Population genetic structure

Shared ancestry between populations was evaluated using ADMIXTURE v 1.22 and the number of cluster that better represent the data was established by crossvalidation as described in^40^. Each ADMIXTURE run was replicated 5 time using different random seeds. Principal Component Analysis (PCA) analysis was performed using EIGENSOFT ^41^.

### Kinship, identical by descent state and runs of homozygosity calculations

Pedigree-based kinship was calculated using the R package Kinship2^42^ using genealogical records. For VBI, pedigrees were reconstructed using data of births, marriages and deaths from the 16th century onwards, collected from churches and city council archives ^21,24^. Pedigree data collection was approved by the institutional ethical committee of the San Raffaele Hospital in Milan and by the Regione Piemonte. Pedigrees of FVG were reconstructed using data collected during sampling and pedigree data collection was approved by the Ethics Committee of the Burlo Garofolo children hospital in Trieste (FVG). Pedigrees of CAR were reconstructed from private data donated to the project and approval was obtained by the local administration of Carlantino, the Health Service of Foggia Province, Italy, and ethical committee of the IRCCS Burlo-Garofolo of Trieste.

Pedigree-based kinship measures provide the expected value of the amount of genome shared identical by descent, however recombinations and mutation introduce variability around the expected values. Genome-based kinship directly accounts for this variability and provide continuous estimates of kinship rather than pedigree-based point estimates (e.g first-degree kinship vary between 0.25+/variability). Pairwise genomic kinship was calculated using KING^43^ (options: --kinship --ibs). A negative genomic kinship coefficient indicates an unrelated pairs and for practical purpose it is safe to rescale negative kinship values to zero ^43^. All analyses except kinship distributions were performed on individuals related at most to 2nd degree according to pedigree-based kinship.

Segments of identity by descent (IBD) and runs of homozigosity (ROH) were identified using the refined IBD algorithm implemented in Beagle v4.r1274^44^ (parameters used: ibd=true, window=24000, overlap=7000). All variants with uncalled genotypes were removed, leaving 7,502,857 sites for further analyses. After discovery we retained IDB and ROH segments with LOD score >5 as threshold to define a true IBD/ROH segment (default value LOD >= 3). When comparing IBD sharing, ROH data and allele frequencies populations were randomly down-sampled to match the size of the smallest village size (FVG-I, 46 unrelated samples) in order to reduce bias from different sample sizes.

### Effective population size

Recent effective population size for each population was estimated using the fraction of genome shared IBD by pairs of individuals^26^. After computational phasing, IBD sharing was computed as described in^45^. We initially removed all variants with minor allele frequency less than 1%. We used a publicly available genetic map (b37, Web Resources), using linear interpolation to infer the genetic position of variants that were not found in the map. We then used the GERMLINE^46^ software to infer IBD sharing across all pairs of samples, using parameters “-min_m 1 -err_hom 2 -err_het 0 -bits 75 -h_extend -homoz”. We computed the density of IBD sharing per genomic region using windows of 0.5 centimorgans (cM), and selected for downstream analysis regions within 5 standard deviations from the genome-wide average, removing regions of unusual sharing, likely due to artifacts or the presence of underlying structural variation. We further imposed a minimum length of 45cM per genomic region to avoid introducing biases due to boundary effects in very short regions. We obtained 27 regions, for a total of ~2050 cM. We computed the fraction of genome shared by the average pair of haploid individuals in each analysed group through segments of length at least 6 cM. We then used the formula: N = (1-f+sqrt(1-f))/2uf derived in^26^, where *f* is the observed fraction of genome shared IBD and *u* is the minimum segment length (in Morgans) to infer effective population size (assuming a constant size). We report diploid effective sizes. Standard errors were computed using the weighted jackknife method^46^, using the 27 genomic regions. To infer an approximate correspondence between IBD segment cut-off and time, we considered the distribution of IBD segments age in a constant population size. Assuming a generation time of 30 years^47^ a cutoff of 6 cM roughly corresponds to IBD segments originating in common ancestors living 30*(100/6)=500 years before present.

### R_x/y_

The metric R_X/Y_ was calculated implementing the formula described in^2,4^ and ^16^ using the information on derived alleles counts per individual in 46 samples per population. Variance of R_X/Y_ was obtained by block jackknife procedure as described in ^48^. To avoid biases due to lack of calling and imputation we considered only sites called as variable in both X and Y populations (shared sites). Information on the ancestral state of the allele were taken from 1000 Genomes annotations (http://ftp.1000genomes.ebi.ac.uk/vol1/ftp/phase1/analysis_results/supporting/ancestral_alignments/). We used four sets of variants with information on functional consequences from Ensembl database v. 78 (www.ensembl.org): (i) 136,805 variants annotated as intergenic, i.e. “in region containing or overlapping no genes that is bounded on either side by a gene, or bounded by a gene and the end of the chromosome” according to Sequence Ontology definition (S.O., http://www.sequenceontology.org/): (ii) 25,773 missense variants, i.e. “A sequence variant, that changes one or more bases, resulting in a different amino acid sequence but where the length is preserved” ( S.O.); (iii) 23,242 synonymous, i.e. “A sequence variant where there is no resulting change to the encoded amino acid” (S.O.), and (iv) 2,804 loss of function variants described in^29^

### Simulations

We used SLiM^49^ to perform forward-in-time simulation of purging of deleterious recessive alleles in a small size human isolated population. SLiM allows simulations of specific genomic structures with different selective pressures acting on them. We simulated a genomic region of 115 kbp containing fifty gene-like structures representative of the average human gene^50^ each composed of 8 exons of 150 bp surrounded by 2 UTR regions of 550 bp. We set the recombination rate to 1.6e-8 recombinations/bp/generation, the mutation rate to 1.2e-8 mutations/bp/generation, and generation time was set to 25 years. Mutations were assumed to be deleterious ¾ of the time in exons and ½ in UTRs. Six deleterious selection coefficients were tested independently (s = -0.2, -0.1, -0.05, -0.01, -0.005 and -0.001). Deleterious mutations were assumed to be fully recessive, additive, or dominant.

For the demographic model we simulated an isolated population of European ancestry undergoing a founder event of different intensity in terms of number of individuals and length of the bottleneck (see simulation scheme in Figure S7) and then evolving under an specific demography. Three different split times of the isolate from Europe (160, 60 and 20 generations ago), and five demographies were tested: continued bottleneck, serial founder effect, bottleneck plus linear expansion, bottleneck plus instantaneous expansion and bottleneck plus linear reduction). To mimic isolation we assumed no migration between the isolate and the European population after the split. The European population was simulated under the model described in^51^ with a minor modifications: because SLiM does not simulate exponential growth we extrapolated the trajectory of the exponential growth found in Out-of-Africa populations from^51^ and increased population effective size every few generations. Finally, in our model the European population separates from the African as described in^51^. To mimic relatedeness we let inbreeding starts one generation after the split between Europe and the isolate using two inbreeding coefficients: one similar to estimates from empirical data (averages among all isolates, f = 0.002, Table S7) and a second one five times bigger (f = 0.01). To use these inbreeding coefficient in the forward simulator (SLiM), we converted inbreeding coefficient (f) to selfing coefficient (σ) using the formula: 

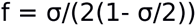
described in (http://darwin.eeb.uconn.edu/eeb348/lecturenotes/inbreeding.pdf).

Each set of parameters and demographies was simulated 500 times and, additionally, we considered groups of 10 replicates together as they were independent chromosomes in order to reduce the variance in the individual recessive load calculation. From every simulated replica we sampled 50 individuals in each of the three populations. In simulations with a final population effective size of less than 50 individuals the whole population was sampled. To speed up computation we used two strategies. First, we removed from simulations the neutrally evolving intergenic and intronic regions, thus mimicking exomic data. Secondly, to avoid repeating the burn-in process, we simulated 500 replicates for Africans and Europeans and saved all the individual genotypes at 200 generations before present (5000 years ago) and we used these population snapshots as genetic pools for the founding nucleus of the isolate while testing different demographic models. We believe that this strategy ensures enough variation for testing properly the different population sizes for the ISO population without having to use a new simulations run from the scratch.

## Acknowledgements

We acknowledge the contribution of people from villages. We also thank Marianna Buonaiuto e Roberto Sirica for helping with the manuscript. This study was supported by the European Commission (FP7/2007–2013, under grant agreement number no. 262055 (ESGI), as a Transnational Access project of the European Sequencing and Genotyping Infrastructure”; Itailan Ministero della Salute - Ricerca Finalizzata PE-2011–02347500 to PG and DT; Italian Ministero della Salute - Public Health Genomics 2010 and Fondazione Cariplo to DT; the Wellcome Trust (Grant Codes WT098051 and WT091310), the NIHR and the EU FP7 (EPIGENESYS Grant Code 257082 and BLUEPRINT Grant Code HEALTH-F5–2011–282510) to NS.

## Author contributions

MC, MP, PFP, EG and VC designed and performed analyses. MT, CFS, SU performed genotype quality controls. YM, RD generated and analysed exome data. AKK coordinated exome sequencing/data generation. PG, DT discussed the analyses' results. NS and VC wrote the manuscript.

## Competing financial interests

The authors declare no competing financial interests.

## References

1. Kimura, M. Population Genetics, Molecular Evolution, and the Neutral Theory. (University of Chicago Press, 1995). at <http://www.press.uchicago.edu/ucp/books/book/chicago/P/bo3645416.html>

2. Do, R. et al. No evidence that selection has been less effective at removing deleterious mutations in Europeans than in Africans. Nat. Genet. 47, 126–131 (2015).

3. Simons, Y. B., Turchin, M. C., Pritchard, J. K. & Sella, G. The deleterious mutation load is insensitive to recent population history. Nat. Genet. 46, 220–224 (2014).

4. Balick, D. J., Do, R., Cassa, C. A., Reich, D. & Sunyaev, S. R. Dominance of Deleterious Alleles Controls the Response to a Population Bottleneck. PLoS Genet. 11, e1005436 (2015).

5. Charlesworth, D. & Willis, J. H. The genetics of inbreeding depression. Nat. Rev. Genet. 10, 783–796 (2009).

6. Whitlock, M. C. Selection, load and inbreeding depression in a large metapopulation. Genetics 160, 1191–1202 (2002).

7. García-Dorado, A. A simple method to account for natural selection when predicting inbreeding depression. Genetics 180, 1559–1566 (2008).

8. Kirkpatrick, null & Jarne, null. The Effects of a Bottleneck on Inbreeding Depression and the Genetic Load. Am. Nat. 155, 154–167 (2000).

9. Glémin, S. How are deleterious mutations purged? Drift versus nonrandom mating. Evol. Int. J. Org. Evol. 57, 2678–2687 (2003).

10. Bittles, A. H. & Neel, J. V. The costs of human inbreeding and their implications for variations at the DNA level. Nat. Genet. 8, 117–121 (1994).

11. Jorde, L. B. Consanguinity and prereproductive mortality in the Utah Mormon population. Hum. Hered. 52, 61–65 (2001).

12. Ober, C., Hyslop, T. & Hauck, W. W. Inbreeding effects on fertility in humans: evidence for reproductive compensation. Am. J. Hum. Genet. 64, 225–231 (1999).

13. Postma, E., Martini, L. & Martini, P. Inbred women in a small and isolated Swiss village have fewer children. J. Evol. Biol. 23, 1468–1474 (2010).

14. Rudan, I. et al. Inbreeding and risk of late onset complex disease. J. Med. Genet. 40, 925–932 (2003).

15. Facon, B. et al. Inbreeding depression is purged in the invasive insect Harmonia axyridis. Curr. Biol. CB 21, 424–427 (2011).

16. Xue, Y. et al. Mountain gorilla genomes reveal the impact of long-term population decline and inbreeding. Science 348, 242–245 (2015).

17. Bittles, A. H., Mason, W. M., Greene, J. & Rao, N. A. Reproductive behavior and health in consanguineous marriages. Science 252, 789–794 (1991).

18. Helgason, A., Pálsson, S., Gudbjartsson, D. F., Kristjánsson, T. & Stefánsson, K. An association between the kinship and fertility of human couples. Science 319, 813–816 (2008).

19. Colonna, V. et al. Small effective population size and genetic homogeneity in the Val Borbera isolate. Eur. J. Hum. Genet. 21, 89–94 (2013).

20. Esko, T. et al. Genetic characterization of northeastern Italian population isolates in the context of broader European genetic diversity. Eur. J. Hum. Genet. 21, 659–665 (2013).

21. Traglia, M. et al. Heritability and demographic analyses in the large isolated population of Val Borbera suggest advantages in mapping complex traits genes. PloS One 4, e7554 (2009).

22. Consortium, T. 1000 G. P. A map of human genome variation from population-scale sequencing. Nature 467, 1061–1073 (2010).

23. 1000 Genomes Project Consortium et al. An integrated map of genetic variation from 1,092 human genomes. Nature 491, 56–65 (2012).

24. Milani, G. et al. Computer-based genealogy reconstruction in founder populations. J. Biomed. Inform. 44, 997–1003 (2011).

25. Pemberton, T. J. et al. Genomic patterns of homozygosity in worldwide human populations. Am. J. Hum. Genet. 91, 275–292 (2012).

26. Palamara, P. F., Lencz, T., Darvasi, A. & Pe'er, I. Length Distributions of Identity by Descent Reveal Fine-Scale Demographic History. Am. J. Hum. Genet. 91, 809–822 (2012).

27. Keinan, A. & Clark, A. G. Recent Explosive Human Population Growth Has Resulted in an Excess of Rare Genetic Variants. Science 336, 740–743 (2012).

28. Coventry, A. et al. Deep resequencing reveals excess rare recent variants consistent with explosive population growth. Nat. Commun. 1, 131 (2010).

29. MacArthur, D. G. et al. A systematic survey of loss-of-function variants in human protein-coding genes. Science 335, 823–828 (2012).

30. Lawrie, D. S., Messer, P. W., Hershberg, R. & Petrov, D. A. Strong purifying selection at synonymous sites in D. melanogaster. PLoS Genet. 9, e1003527 (2013).

31. Li, H. et al. The Sequence Alignment/Map format and SAMtools. Bioinforma. Oxf. Engl. 25, 2078–2079 (2009).

32. DePristo, M. A. et al. A framework for variation discovery and genotyping using next-generation DNA sequencing data. Nat. Genet. 43, 491–498 (2011).

33. Browning, S. R. & Browning, B. L. Rapid and Accurate Haplotype Phasing and Missing-Data Inference for Whole-Genome Association Studies By Use of Localized Haplotype Clustering. Am. J. Hum. Genet. 81, 1084–1097 (2007).

34. Delaneau, O., Howie, B., Cox, A. J., Zagury, J.-F. & Marchini, J. Haplotype Estimation Using Sequencing Reads. Am. J. Hum. Genet. 93, 687–696 (2013).

35. Howie, B. N., Donnelly, P. & Marchini, J. A Flexible and Accurate Genotype Imputation Method for the Next Generation of Genome-Wide Association Studies. PLoS Genet 5, e1000529 (2009).

36. Howie, B., Marchini, J. & Stephens, M. Genotype Imputation with Thousands of Genomes. G3 Genes Genomes Genet. 1, 457–470 (2011).

37. Sherry, S. T. et al. dbSNP: the NCBI database of genetic variation. Nucleic Acids Res. 29, 308–311 (2001).

38. McLaren, W. et al. Deriving the consequences of genomic variants with the Ensembl API and SNP Effect Predictor. Bioinforma. Oxf. Engl. 26, 2069–2070 (2010).

39. Wang, Y., Lu, J., Yu, J., Gibbs, R. A. & Yu, F. An integrative variant analysis pipeline for accurate genotype/haplotype inference in population NGS data. Genome Res. 23, 833–842 (2013).

40. Alexander, D. H., Novembre, J. & Lange, K. Fast model-based estimation of ancestry in unrelated individuals. Genome Res. 19, 1655–1664 (2009).

41. Patterson, N., Price, A. L. & Reich, D. Population structure and eigenanalysis. PLoS Genet. 2, e190 (2006).

42. Sinnwell, J. P., Therneau, T. M. & Schaid, D. J. The kinship2 R package for pedigree data. Hum. Hered. 78, 91–93 (2014).

43. Manichaikul, A. et al. Robust relationship inference in genome-wide association studies. Bioinformatics 26, 2867–2873 (2010).

44. Browning, B. L. & Browning, S. R. Improving the Accuracy and Efficiency of Identity-by-Descent Detection in Population Data. Genetics 194, 459–471 (2013).

45. The Genome of the Netherlands Consortium. Whole-genome sequence variation, population structure and demographic history of the Dutch population. Nat. Genet. 46, 818–825 (2014).

46. Gusev, A. et al. Whole population, genome-wide mapping of hidden relatedness. Genome Res. 19, 318–326 (2009).

47. Fenner, J. N. Cross-cultural estimation of the human generation interval for use in genetics-based population divergence studies. Am. J. Phys. Anthropol. 128, 415–423 (2005).

48. Busing, F. M. T. A., Meijer, E. & Leeden, R. V. D. Delete-m Jackknife for Unequal m. Stat. Comput. 9, 3–8 (1999).

49. Messer, P. W. SLiM: simulating evolution with selection and linkage. Genetics 194, 1037–1039 (2013).

50. Sakharkar, M. K., Chow, V. T. K. & Kangueane, P. Distributions of exons and introns in the human genome. In Silico Biol. 4, 387–393 (2004).

51. Gravel, S. et al. Demographic history and rare allele sharing among human populations. Proc. Natl. Acad. Sci. U. S. A. 108, 11983–11988 (2011). 415–423 (2005).

